# Perceived facial happiness during conversation correlates with insular and hypothalamus activity for humans, not robots: An investigation of the somatic marker hypothesis applied to social interactions

**DOI:** 10.1101/2021.11.22.469413

**Authors:** Thierry Chaminade, Nicolas Spatola

## Abstract

The somatic marker hypothesis posits that perceiving emotions entails reenacting markers of self emotions, in particular in the autonomous nervous system. Well studied in decision-making tasks, it has not been tested in a social cognitive neuroscience framework, and in particular for the automatic processing of positive emotions during natural interactions. Here, we address this question using a unique corpus of brain activity recorded during unconstrained conversations between participants and a human or a humanoid robot. fMRI recordings are used to test whether activity in the most important brain regions in relation to the autonomic system, the amygdala, hypothalamus and insula, is affected by the level of happiness expressed by the human and robot agents. Results indicate that for the hypothalamus and the insula, in particular the anterior agranular region strongly involved in processing social emotions, activity in the right hemisphere increases with the level of happiness expressed by the human, but not the robot. Results indicate that perceiving positive emotions in social interactions induces the local brain responses predicted by the somatic marker hypothesis, but only when the interacting agent is a real human.

## 1 Introduction

Human beings are intrinsically social creatures whose lives are intertwined with others’. Engaged in any social interaction, we cannot help but gather information about others’ mental states, in particular their emotions that signal the quality of the ongoing interaction. Facial expressions convey many signals on people’s mental states that are extremely relevant for social interactions. Our capacity to take into account these social facial signals makes it possible to adapt our behaviour when relating to others. It is even possible that information carried on our face influences behaviour in the absence of explicit perception, as it has been found for pupil size (Harrison et al., 2009) and suggested for subtle face blushing (Crozier, 2001). Understanding mechanisms involved in processing this dynamic visual information in naturalistic social interactions is required for the advancement of second-person neuroscience (Redcay & Schilbach, 2019), that advocates studies of real-time and reciprocal exchanges between individuals.

As other perceptual mechanisms involved in social interactions such as the perception of faces and voices (Belin, 2017), processing of facial expression of emotions comprises at least two mechanisms (Adolphs, 2002). First, a fast visual mechanism that identifies the geometric configuration of facial features, that can be described as visual norm-based coding of facial patterns of muscular activations (also known as Action Units). The later stage has been proposed to involve embodied representations of emotions, making use of somatic markers of the emotions felt by the individual to evaluate both the valence and the arousal of the perceived emotions (Damasio et al., 1996). This reenactment of felt emotions is believed to be grounded in a distributed network of brain regions devoted to emotion processing (Poppa & Bechara, 2018; Vuilleumier & Pourtois, 2007). Brain regions include a number of cortical as well as various subcortical regions, in contrast to the strictly cortical correlates of sensory processing.

The somatic marker hypothesis postulates that autonomic responses associated with specific emotions are re-enacted in response to certain stimuli (Damasio et al., 1996). At the core of the brain networks involved in associating peripheral and central nervous systems are the interconnected insular cortex and subcortical amygdala and hypothalamus (Damasio et al., 2000; Pessoa, 2017), brain regions that share a number of characteristics. They are ubiquitous in the vertebrate kingdom, they are involved in core body functions such as homeostasis, and they are smaller in humans, compared to other primates, relative to other subcortical (e.g. hippocampus) or cortical (e.g. prefrontal cortex) areas. It could be that, in most vertebrates, emotions have a direct relation with survival thus play an important role in natural selection. In particular fear is associated with specific homeostatic and behavioral “fight-or-flight” responses essential for animal survival.

Neural mechanisms involved in processing of emotions in these regions are not comparable to those described in cortical areas dedicated to sensory perception. The reason is that processing of facial and vocal identity at the cortical level, in the occipital and temporal lobes respectively, are important for face-to-face conversations, for which humans have been specialized in comparison to other animals, even primates. Yet emotions play an important, though not-so-well understood, role in social cognition, and thus deserve to be investigated within second-person cognitive neuroscience framework. Such an investigation is difficult given the incompatibility between the necessary experimental control in experimental research and the freedom associated with natural social behaviours. Exceptional circumstances, such as intracerebral sampling of hormones in narcoleptic patients clearly demonstrates that the amygdala physiology is influenced by social emotions (Blouin et al., 2013). Artificial agents provide an experimental tool to bypass this difficulty. Indeed, human-like robots that can produce facial emotions have been developed, either in mechanical form (Chaminade et al., 2010) or through a projection (Al Moubayed et al., 2012). These agents allowed the development of new paradigms to better understand mechanisms involved in human social interaction (Chaminade, 2017). They are unique tools to dissociate perceptual, bottom-up processes, that are automatic and irresistible (such as speech understanding), from contextual, top-down processes, that are influenced by a large number of factors amongst which the perceiver’s mental states play a major role.

For instance, two fMRI studies comparing brain response to passive viewing of humans’ and robots’ facial expressions of emotions reported reduced activity for the robot expressions of emotions, in the left insula and right amygdala (Hortensius et al., 2018). Another fMRI study (Chaminade et al., 2010) found reduced activity in the hypothalamus when participants believed they were playing chifumi with a robot compared to a human, without seeing their opponent (Chaminade et al., 2015). However, these studies are not sufficient to answer the crucial question: are these effects due to the nature of the agent, a dichotomic factor, or rather to quantitative differences in the intensity of the social emotions conveyed by the interaction?

Here we investigate this question, important both for understanding natural human social cognition and for the future of human-robot interactions, by analyzing a unique fMRI corpus of human-human and human-robot face-to-face conversations which represent social human-robot interactions (Rauchbauer et al., 2019). In contrast to considering “the six emotions” as a category, following a long trend of research initiated by Charles Darwin (Darwin, 1872) and comforted one century later by Paul Ekman (Ekman, 1974), we propose that the main emotion for social bonding is the positive emotion of happiness - facial and vocal expression of happiness convey comfort in the interaction. We automatically extracted facial happiness expressed by the human and robot agent and used it as covariate in multivariate analyses of the activity extracted in functional subregions of the insula, in two divisions of the amygdala, and in the hypothalamus, bilaterally. We expected that, applying the somatic marker hypothesis to positive emotions perceived by the participants during a natural conversation, response in these particular brain areas would be correlated with the level of happiness expressed by the human, but not the robot.

## 2 Methods

As the analysis presented here is based on an existing shared corpus described *in extenso* in previous publications (Rauchbauer et al., 2019, 2020), we will only summarize the information necessary to support the results and discussion but readers are referred to the technical publications for more details.

### 2.1 Data acquisition

When participants (n = 24, 17 women, μ = 26.76 years, σ = 7.96) arrived at the MRI center, one experimenter presented them the cover story, that they would participate to a neuromarketing experiment by discussing online about the meaning of images purportedly prepared for two upcoming advertising campaigns. The experimenter then introduced them to the agents they would be discussing with. The human partner was a confederate of the experimenter gender-matched with each participant, and the artificial agent was a back-projected conversational robotic head (Al Moubayed et al., 2012) with appearance, voice and accessories reminiscent of the natural agent (Figure 1).

**Figure 1.**
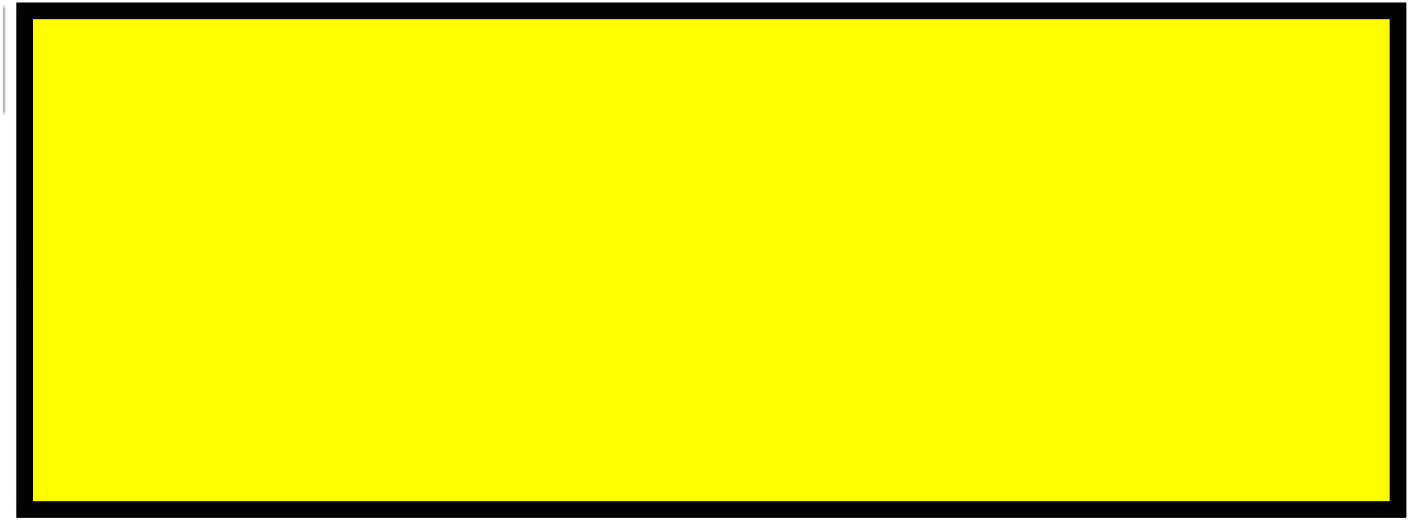
Snapshots from the live video feeds projected to the scanned participants, with the Human (left) and Robot (right) interlocutor.

During functional magnetic resonance imaging (fMRI) acquisition, participants discussed online with these two agents alternatively while lying supine in the scanner and having their brain activity recorded. BOLD signal (Blood Oxygen-Level Dependent) was recorded in four sessions of fMRI acquisition, each lasting approximately eight minutes. Each session comprised 6 experimental trials that proceeded as follows: a picture appeared for 8.3 seconds, then after a 3 seconds pause with a white fixation cross on a black background, a one-minute live conversation took place with either the human or the artificial agent, alternatively. The participant and interlocutor could hear each other in real-time, and the participant additionally saw a life feed of the interlocutor. All audios and videos available were recorded for further analysis. In total, each participant took part in twelve one-minute conversations with the human and twelve one-minute conversations with the artificial agent. NOte that other variables were recorded, such as the participants’ eye movements, but won’t be discussed further here since they are not used in the analysis presented here.

Unbeknown to the participants who believed the robot was autonomous, it was actually controlled remotely by the confederate acting as the human interlocutor. A set of pre-recorded answers, chosen following a previous version of the experiment, were selected in real-time by pressing virtual buttons on a touch-sensitive tablet. Examples of answers, some generic (e.g. “yes”, “no”, “maybe”, “I don’t know”) and others specific to an image (“It’s a yellow pear”) or to one of the two advertising campaigns (“Maybe it’s a campaign to promote local fruits cultivation”) are given in Table 1. Responses were stored as texts and played using the voice synthesizer provided with the robot.

### 2.2 Data preparation

The processing of fMRI data followed standard procedures. The volumes acquired represent the BOLD signal in 2.5 x 2.5 x 2.5 cm^3^ voxels of the brain. Each volume contains 54 slices of 84 x 84 in-plane voxel, and is recorded in 1.205 seconds. Preprocessing entails a correction of temporal synchronization of the acquired slices, a realignment of the volumes of each session on the first one, and a correction of the deformations due to the local distortions of the magnetic field and participants’ movements. Normalization uses the DARTEL procedure (Ashburner, 2007) to put the imaged brains of all participants in the standard MNI space. Several nuisance covariates were computed to eliminate motion artifacts, potential blood pulse and respiration artifacts, which were highly relevant in a paradigm involving speech, as well as global grey matter signal, white matter activity, and cerebrospinal fluid activity to control global signal fluctuations unrelated to the task (TAPAS toolkit, Kasper et al., 2017).

The analysis of fMRI data was first based on the general linear model implemented in SPM (Friston et al., 2007). Each trial was modelled as a single regressor, and the images presented before each discussion were modelled as a single regressor. We used a brain parcellation formed from functional and connectivity brain data, the Brainnetome atlas (Fan et al., 2016), so that the 246 regions of interest of the atlas represent sets of voxels that are homogeneous in terms of function. For each of the 24 trials (12 for the human, 12 for the robot) of each of the 24 participants, we extracted the average response across the one-minute duration using the MarsBAR toolbox (Brett et al., 2002). We then focused on regions of interest (ROIs) that belong to the insular and subcortical insular regions that form the core of the somatic emotional system. The insula can be parcellated according to a gradient of increasing granularity from front to back. Within the Brainnetome atlas numbering systems, insular regions are (in left/right hemispheres) 165/166 (ventral Agranular, [vA]) and 167/168 (dorsal Agranular, [dA]) anteriorly, 169/170 (ventral Dysgranular, [vD]) and 173/174 (dorsal Dysgranular, [dD]) in intermediate location, and posteriorly 171/172, (dorsal Granular insula, [dG]) and 163/164 (Hypergranular insula [H]). The amygdala is composed of regions 211/212 (Medial part of the amygdala [M], roughly corresponding to the medial and basal nuclei) and 213/214 (Lateral part of the amygdala [L], roughly corresponding to central and lateral nuclei; Saygin et al., 2011). Activity of the hypothalamus [Hy] was also extracted using a mask previously developed (Wolfe et al., 2015) given that this key region for homeostasis is not featured in the Brainnetome atlas. Figure 2 can be used to visualize all regions.

**Fig 2.**
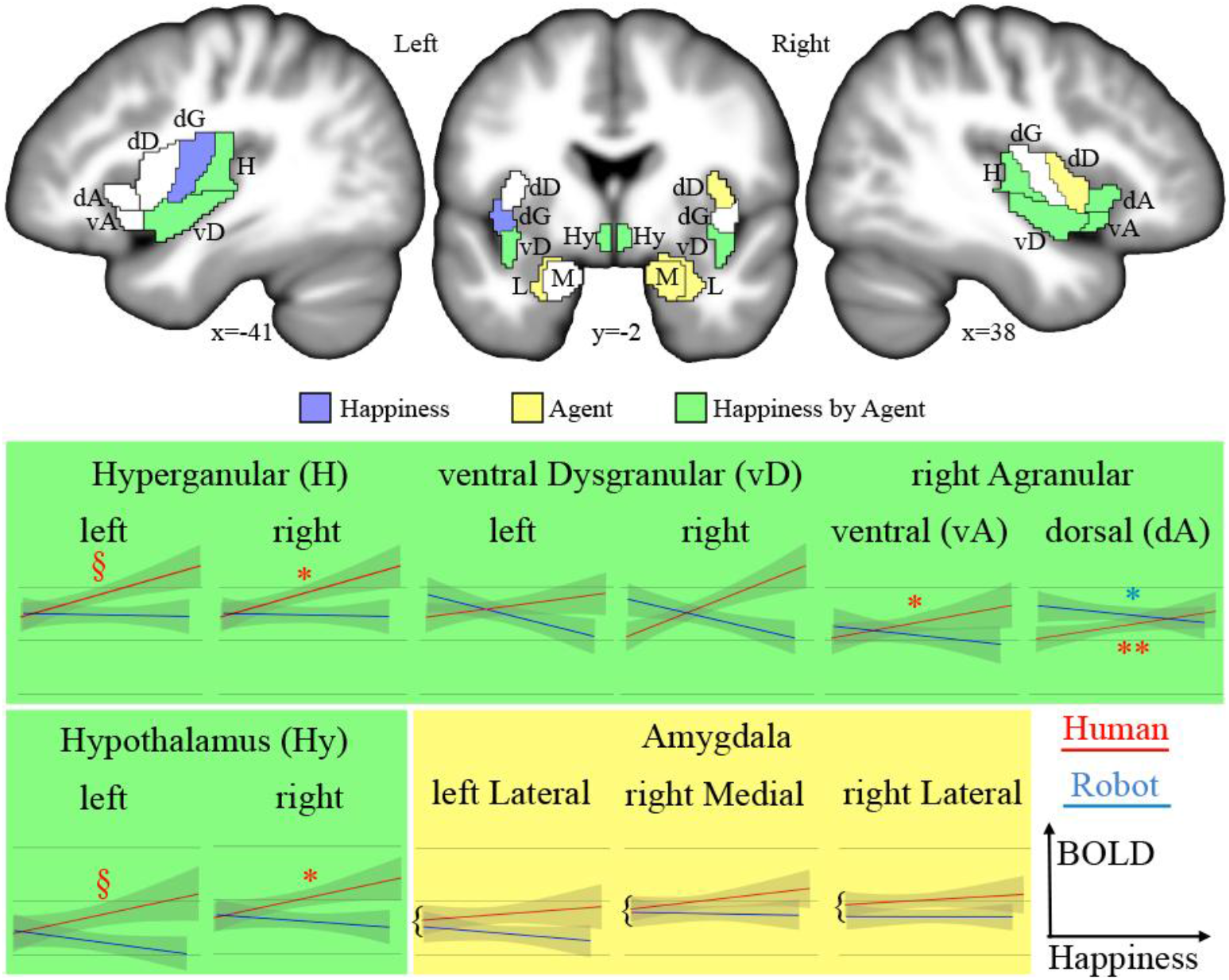
Top: regions of interests (acronyms described in Table 2) overlaid on sections of the average of participants’ brains after normalization. In blue are the areas showing a significant main effect of Happiness, in yellow a significant main effect of the type of Agent and in green a significant interaction. Bottom: plots of significant effects on BOLD response (arbitrary units [AU] after normalization) against the level of Happiness expression for human (red) and robot (blue) in ROIs discussed in the text. The significant correlations are indicated **: *p* < 0. 001, *:*p* < 0.050, §:*p* < 0.100.

### 2.3 Data analysis

Facial emotions were extracted automatically from the videos of the human and robot interlocutors using a freely available machine learning algorithm (Arriaga et al., 2019) available at https://github.com/oarriaga/face_classification. The algorithm uses a Convolutional Neural Network to classify facial expression within 7 classes (Anger, Disgust, Fear, Happiness, Sadness, Surprise, and Neutral), and achieves an accuracy of 66% on a dataset of manually labelled 35887 grayscale images. The recordings used for this classification, with 640 by 480 images of full-screen faces of the human or robot interlocutor looking directly on the screen, were appropriate for this automatic classification. The probability predicted for each emotion was extracted frame by frame, then summed over all frames of a trial, therefore providing one score for each emotion per trial.

Statistical analyses were performed in R with the package lme4. Multivariate models were used as the effect of each predictor (level of facial happiness, type of agent) is evaluated while holding constant the effect of the other predictors on the dependent variable. We introduced the happiness score obtained for each trial as a predictive continuous variable for all regions evaluated, with the nature of the agent the only categorical factor of interest. Other factors (identity of the participant, session number) were used as random variables. The models were selected in a backward process based on the restricted maximum likelihood.

## 3 Results

There is a significant difference between the Happiness extracted for the human and robot interlocutors (*t* = 21.08, *p* < 0.001, CI_95%_ [0.89, 2.76]), with higher scores for the human agent. This significant difference argues in favor of the multivariate linear model approach that allows to evaluate independently each term’s unique contribution to the variation of the BOLD signal, therefore controlling for the significant difference of Happiness between agents.

We tested the effect of the nature of the dichotomous factor describing the nature, human or robotic, of the participant’s interlocutor (Agent) and of the continuous happiness score (Happiness), as well as the interaction between these terms (Happiness by Agent) on the BOLD signal in the insula, amygdala and hypothalamus ROIs. Statistical results for all ROIs are presented in Table 2 and illustrated on Figure 2 (top). The interaction term is particularly important, as it indicates that the happiness score affects local brain response differently depending on whether the interlocutor is the human or the robot. This term reveals different but consistent patterns in the amygdala ROIs on the one hand, for which it is never significant, and in the insula and hypothalamus ROIs on the other hand, for which it is significant in many regions.

**Table 2.**
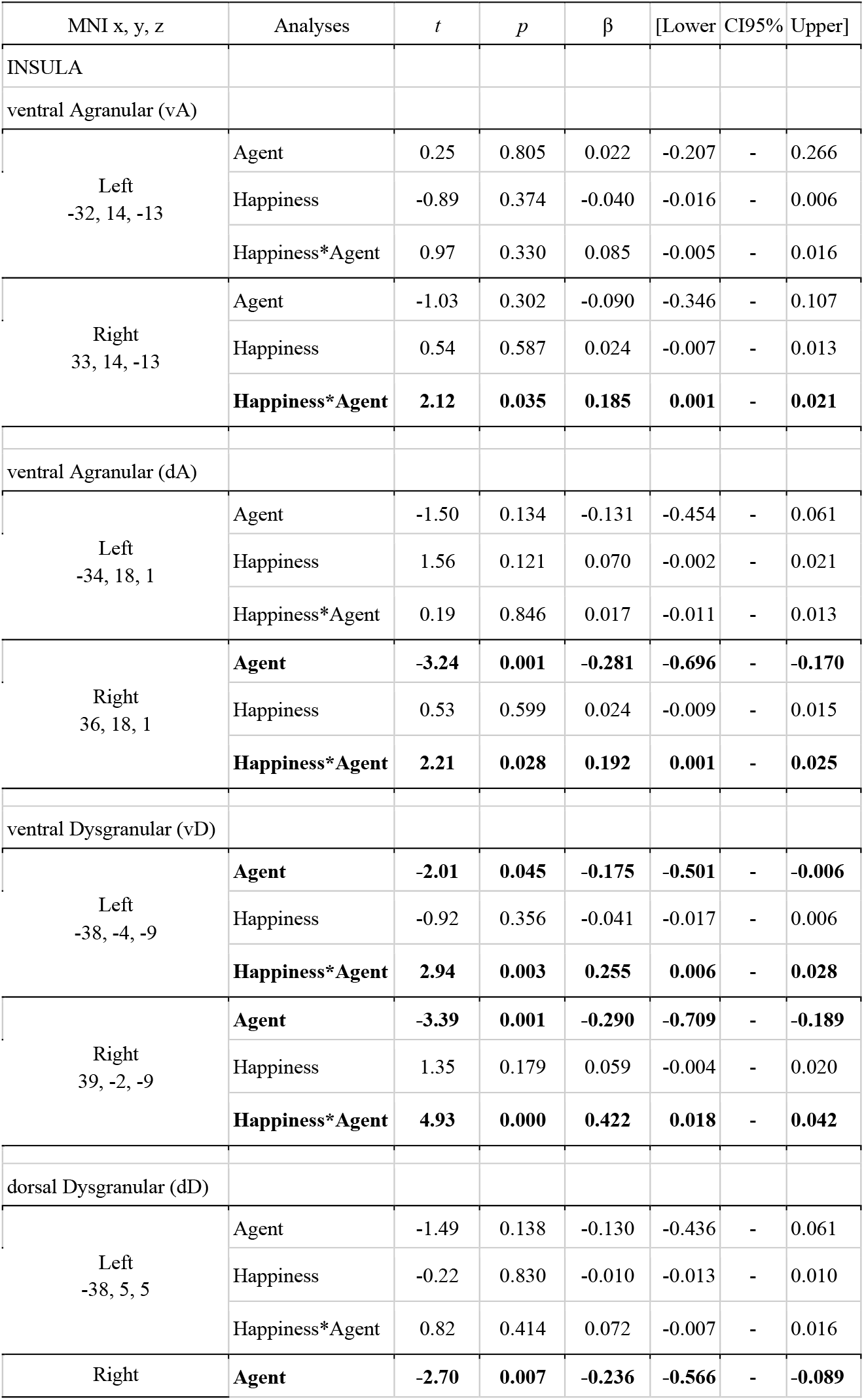
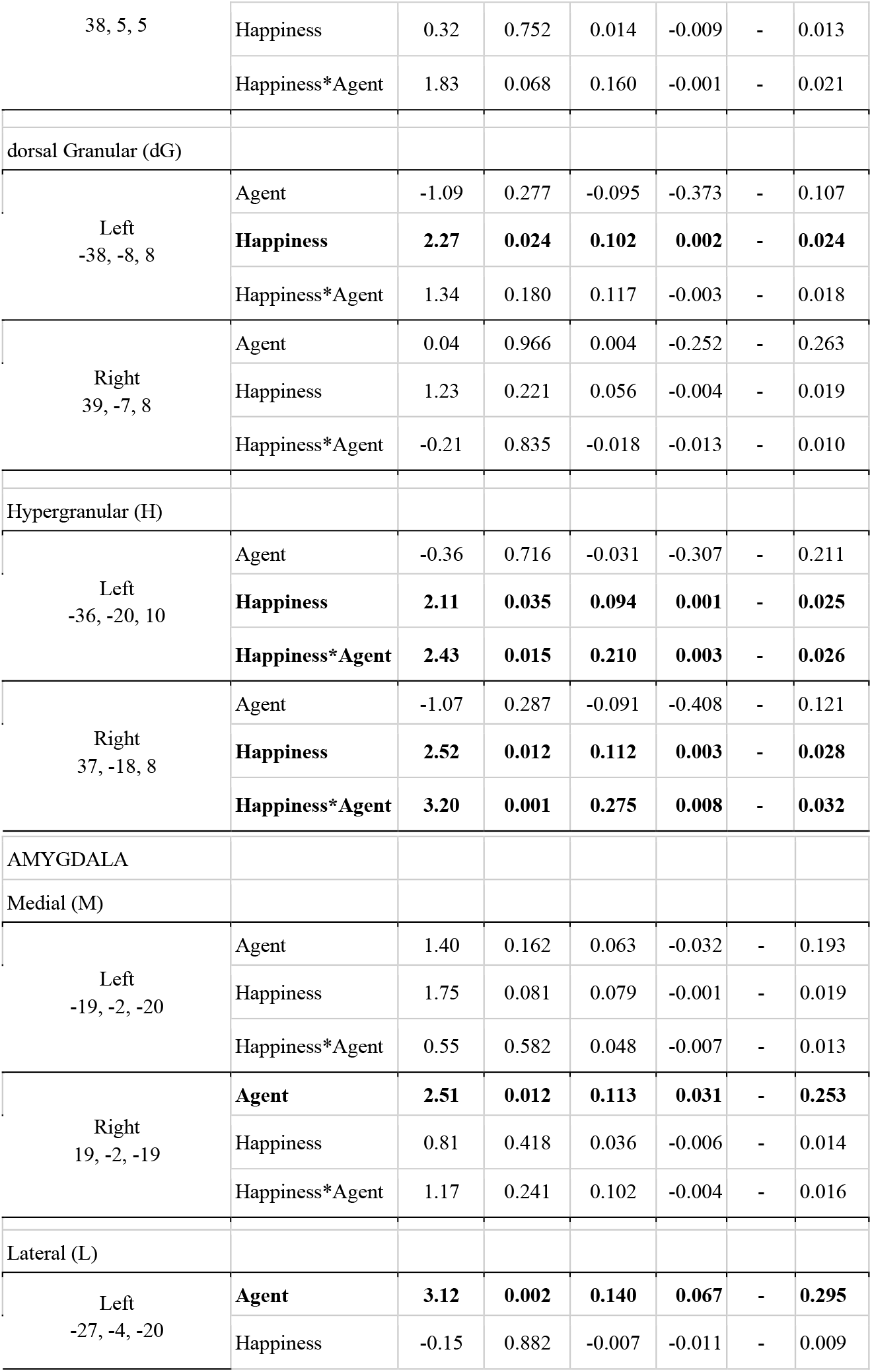
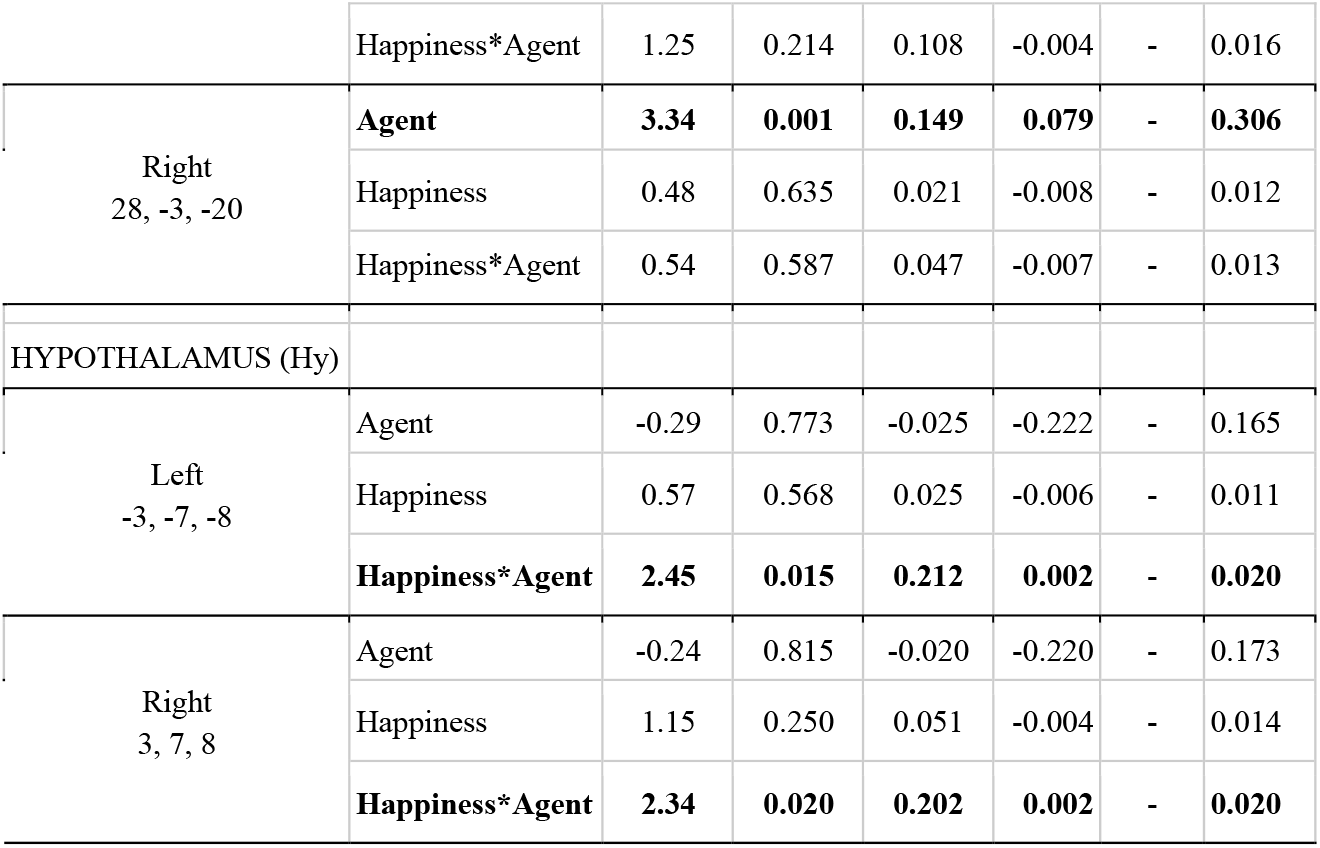
Results of the statistical analysis. Significant effects (*p* < 0.05) are indicated in bold.

Brain response to Natural and Artificial Agents' Happiness
| MNI x, y, z | Analyses | *t* | *p* | $\beta$ | [Lower | CI95% | Upper] |
| --- | --- | --- | --- | --- | --- | --- | --- |
| INSULA |  |  |  |  |  |  |  |
| ventral Agranular (vA) |  |  |  |  |  |  |  |
| Left<br>-32, 14, -13 | Agent | 0.25 | 0.805 | 0.022 | -0.207 | - | 0.266 |
|  | Happiness | -0.89 | 0.374 | -0.040 | -0.016 | - | 0.006 |
|  | Happiness*Agent | 0.97 | 0.330 | 0.085 | -0.005 | - | 0.016 |
| Right<br>33, 14, -13 | Agent | -1.03 | 0.302 | -0.090 | -0.346 | - | 0.107 |
|  | Happiness | 0.54 | 0.587 | 0.024 | -0.007 | - | 0.013 |
|  | **Happiness\*Agent** | **2.12** | **0.035** | **0.185** | **0.001** | - | **0.021** |
| ventral Agranular (dA) |  |  |  |  |  |  |  |
| Left<br>-34, 18, 1 | Agent | -1.50 | 0.134 | -0.131 | -0.454 | - | 0.061 |
|  | Happiness | 1.56 | 0.121 | 0.070 | -0.002 | - | 0.021 |
|  | Happiness*Agent | 0.19 | 0.846 | 0.017 | -0.011 | - | 0.013 |
| Right<br>36, 18, 1 | **Agent** | **-3.24** | **0.001** | **-0.281** | **-0.696** | - | **-0.170** |
|  | Happiness | 0.53 | 0.599 | 0.024 | -0.009 | - | 0.015 |
|  | **Happiness\*Agent** | **2.21** | **0.028** | **0.192** | **0.001** | - | **0.025** |
| ventral Dysgranular (vD) |  |  |  |  |  |  |  |
| Left<br>-38, -4, -9 | **Agent** | **-2.01** | **0.045** | **-0.175** | **-0.501** | - | **-0.006** |
|  | Happiness | -0.92 | 0.356 | -0.041 | -0.017 | - | 0.006 |
|  | **Happiness\*Agent** | **2.94** | **0.003** | **0.255** | **0.006** | - | **0.028** |
| Right<br>39, -2, -9 | **Agent** | **-3.39** | **0.001** | **-0.290** | **-0.709** | - | **-0.189** |
|  | Happiness | 1.35 | 0.179 | 0.059 | -0.004 | - | 0.020 |
|  | **Happiness\*Agent** | **4.93** | **0.000** | **0.422** | **0.018** | - | **0.042** |
| dorsal Dysgranular (dD) |  |  |  |  |  |  |  |
| Left<br>-38, 5, 5 | Agent | -1.49 | 0.138 | -0.130 | -0.436 | - | 0.061 |
|  | Happiness | -0.22 | 0.830 | -0.010 | -0.013 | - | 0.010 |
|  | Happiness*Agent | 0.82 | 0.414 | 0.072 | -0.007 | - | 0.016 |
| Right | **Agent** | **-2.70** | **0.007** | **-0.236** | **-0.566** | - | **-0.089** |
| 38, 5, 5 | Happiness | 0.32 | 0.752 | 0.014 | -0.009 | - | 0.013 |
|  | Happiness*Agent | 1.83 | 0.068 | 0.160 | -0.001 | - | 0.021 |
| dorsal Granular (dG) |  |  |  |  |  |  |  |
| Left<br>-38, -8, 8 | Agent | -1.09 | 0.277 | -0.095 | -0.373 | - | 0.107 |
|  | **Happiness** | **2.27** | **0.024** | **0.102** | **0.002** | - | **0.024** |
|  | Happiness*Agent | 1.34 | 0.180 | 0.117 | -0.003 | - | 0.018 |
| Right<br>39, -7, 8 | Agent | 0.04 | 0.966 | 0.004 | -0.252 | - | 0.263 |
|  | Happiness | 1.23 | 0.221 | 0.056 | -0.004 | - | 0.019 |
|  | Happiness*Agent | -0.21 | 0.835 | -0.018 | -0.013 | - | 0.010 |
| Hypergranular (H) |  |  |  |  |  |  |  |
| Left<br>-36, -20, 10 | Agent | -0.36 | 0.716 | -0.031 | -0.307 | - | 0.211 |
|  | **Happiness** | **2.11** | **0.035** | **0.094** | **0.001** | - | **0.025** |
|  | **Happiness\*Agent** | **2.43** | **0.015** | **0.210** | **0.003** | - | **0.026** |
| Right<br>37, -18, 8 | Agent | -1.07 | 0.287 | -0.091 | -0.408 | - | 0.121 |
|  | **Happiness** | **2.52** | **0.012** | **0.112** | **0.003** | - | **0.028** |
|  | **Happiness\*Agent** | **3.20** | **0.001** | **0.275** | **0.008** | - | **0.032** |
| AMYGDALA |  |  |  |  |  |  |  |
| Medial (M) |  |  |  |  |  |  |  |
| Left<br>-19, -2, -20 | Agent | 1.40 | 0.162 | 0.063 | -0.032 | - | 0.193 |
|  | Happiness | 1.75 | 0.081 | 0.079 | -0.001 | - | 0.019 |
|  | Happiness*Agent | 0.55 | 0.582 | 0.048 | -0.007 | - | 0.013 |
| Right<br>19, -2, -19 | **Agent** | **2.51** | **0.012** | **0.113** | **0.031** | - | **0.253** |
|  | Happiness | 0.81 | 0.418 | 0.036 | -0.006 | - | 0.014 |
|  | Happiness*Agent | 1.17 | 0.241 | 0.102 | -0.004 | - | 0.016 |
| Lateral (L) |  |  |  |  |  |  |  |
| Left<br>-27, -4, -20 | **Agent** | **3.12** | **0.002** | **0.140** | **0.067** | - | **0.295** |
|  | Happiness | -0.15 | 0.882 | -0.007 | -0.011 | - | 0.009 |
|  | Happiness*Agent | 1.25 | 0.214 | 0.108 | -0.004 | - | 0.016 |
| Right<br>28, -3, -20 | **Agent** | **3.34** | **0.001** | **0.149** | **0.079** | - | **0.306** |
|  | Happiness | 0.48 | 0.635 | 0.021 | -0.008 | - | 0.012 |
|  | Happiness*Agent | 0.54 | 0.587 | 0.047 | -0.007 | - | 0.013 |
| HYPOTHALAMUS (Hy) |  |  |  |  |  |  |  |
| Left<br>-3, -7, -8 | Agent | -0.29 | 0.773 | -0.025 | -0.222 | - | 0.165 |
|  | Happiness | 0.57 | 0.568 | 0.025 | -0.006 | - | 0.011 |
|  | **Happiness\*Agent** | **2.45** | **0.015** | **0.212** | **0.002** | - | **0.020** |
| Right<br>3, 7, 8 | Agent | -0.24 | 0.815 | -0.020 | -0.220 | - | 0.173 |
|  | Happiness | 1.15 | 0.250 | 0.051 | -0.004 | - | 0.014 |
|  | **Happiness\*Agent** | **2.34** | **0.020** | **0.202** | **0.002** | - | **0.020** |

Results provided in Table 3 and illustrated on Figure 2 (bottom) indicate significant positive correlation between BOLD signal and Happiness for the human agent in two right hemisphere insula ROIs (dorsal Agranular and Hypergranular) as well as a trend for the left Hypergranular region. In parallel, the analyses revealed negative correlation for the robot agent in the right ventral and dorsal Agranular insula ROIs. The correlation was positive for the human Agent in the hypothalamus (significant in right; trend in left) and not significant for the robot.

**Table 3.** Effect of Happiness on BOLD signal calculated separately for the human and robot agent in ROIs having a significant Agent * Happiness interaction. Significant correlations (*p* < 0.050) are indicated in bold.

|  | Human |  |  |  |  |  | Robot |  |  |  |  |  |
| --- | --- | --- | --- | --- | --- | --- | --- | --- | --- | --- | --- | --- |
| Regions | *t* | *p* | $\beta$ | [Lower | CI95% | Upper] | *t* | *p* | $\beta$ | [Lower | CI95% | Upper] |
| INSULA |  |  |  |  |  |  |  |  |  |  |  |  |
| ventral Agranular (vA), right | 1.46 | 0.147 | 0.092 | -0.004 | - | 0.027 | **-2.68** | **0.008** | **-0.167** | **-0.038** | - | **-0.006** |
| dorsal Agranular (dA), right | **4.34** | **< 0.001** | **0.265** | **0.021** | - | **0.055** | **-2.59** | **0.010** | **-0.160** | **-0.038** | - | **-0.005** |
| ventral Dysgranular (vD), left | 1.55 | 0.121 | 0.098 | -0.003 | - | 0.027 | -1.05 | 0.295 | -0.066 | -0.024 | - | 0.007 |
| ventral Dysgranular (vD), right | 0.43 | 0.667 | 0.027 | -0.012 | - | 0.019 | -0.72 | 0.470 | -0.046 | -0.022 | - | 0.010 |
| Hypergranular (H), left | 1.90 | 0.059 | 0.119 | -0.001 | - | 0.028 | -1.10 | 0.271 | -0.070 | -0.023 | - | 0.006 |
| Hypergranular (H), right | **2.02** | **0.044** | **0.127** | **0.000** | - | **0.033** | -1.15 | 0.252 | -0.072 | -0.028 | - | 0.007 |
| HYPOTHALAMUS (Hy), left | 1.93 | 0.054 | 0.121 | 0.000 | - | 0.027 | -1.50 | 0.136 | -0.094 | -0.020 | - | 0.003 |
| HYPOTHALAMUS (Hy), right | **2.41** | **0.017** | **0.150** | **0.003** | - | **0.029** | -0.86 | 0.391 | -0.054 | -0.018 | - | 0.007 |

## 4 Discussion

The robotic implementation used in this experiment had clear limitations in the expression of emotion, both visually and orally. its happiness was restricted to smiling. the human behaved naturally, which included, for example, occasional laughter. Consequently, humans had quantitatively higher levels of Happiness than the robot. As was expected given these limitations, we found a significant increase in the level of happiness exhibited by the human compared to the robot. The potential statistical bias that could result from this intrinsic difference was controlled in the rest of the analyses by using multivariate models.

We used brain activity (BOLD signal) recorded with fMRI when participants discussed freely with a human or a robot agent to investigate how emotions are processed in the brain. More precisely, we wanted to test whether brain areas involved in sensing and/or controlling body homeostasis, believed to play an important role in the feeling and perception of emotions according to the somatic marker hypothesis (Damasio et al., 1996), respond differently when the emotion is carried by a human compared to a robot. Results indicate different profiles of response depending on the region being investigated. But before getting into the details, general remarks can be made. First, there are statistically significant results, demonstrating that despite the difficulties related to the approach chosen, that is, recording brain activity during unconstrained behaviours and relating the activity afterwards to conversational characteristics extracted from the recorded behaviours, here the happiness expressed by the human and robot locutor, this analysis allows to reveal brain correlates associated with natural interactions. This is particularly important as such approaches should become widespread in second-person neuroscience. Second, in all of the regions investigated, activity is significantly affected by the experimental terms, even if sometimes only in hemisphere, in support of the right choice of regions to investigate perception of emotions. Finally, the finding that not only the factor describing the nature of the interlocutor, human or robot, but also its interaction term crossing the more subtle happiness score estimated automatically, yield significant results is in direct agreement with one central assumption of this project, that during natural social interactions, the nature of the interacting agent, natural or artificial, is an outstanding factor influencing the behavior and associated neural correlates. Therefore, in addition to providing insights into the brain correlates of natural social interactions, these results could also be crucial to better understand human behaviour in human-robot interactions and ultimately contribute to improving the social competence of humanoid robots.

In the amygdala, only the main effect of the factor Agent was statistically significant, with higher BOLD signal during human, compared to robot, trials in 3 out of the 4 ROIs (lateral bilaterally, medial left hemisphere). As explained in the introduction, emotions are a complex construct quite difficult to investigate, and in animal models, up to now, fear has been the focus of most attention thanks to the paradigm of fear conditioning, that allows investigating this emotion without requiring verbal assessment of the participant’s feeling. Thanks to animal studies, it is well known that the amygdala is necessary for fear conditioning, and takes the form of long-term potentiation within fear circuits. It is interesting that the lateral parts of the amygdala depict a larger effect of agent than the medial division, as the latter is more associated with visceral inputs compared to the lateral being more associated with sensory, including visual and auditory, inputs (LeDoux, 2007). The Agent effect could thus be due to the large difference in the aspect and voice of the human and robot agents, but the larger response for the robot is not compatible with this interpretation as the robot had less expressions and speech than the human. Alternatively, it could be related to a repulsion caused by the robot postulated by the uncanny valley hypothesis (Mori et al., 2012). Importantly, we don’t reproduce the modulation of amygdalar activity by the type of social interaction reported using a totally different methodology (Blouin et al., 2013).

In the insula and the hypothalamus, the happiness displayed affected the brain activity differently depending on the nature of the agent. In regions with significant interactions, plotting the BOLD response as a function of Happiness, shown on green background in Figure 2, bottom, always resulted in a similar trend, with an increase of response for the human agent and no effect or a decrease for the human agent. Increases for the human agent were statistically significant in the right hemisphere for the hypothalamus, the hypergranular and dorsal agranular insula region, and the decrease for the robot was significant for the right dorsal agranular insula region.

Insula and hypothalamus are closely related to embodied emotional processing and are known to be influenced by social context (Bartz et al., 2011; Bernhardt & Singer, 2012). The functions of these regions can be described by the somatic markers hypothesis, the former receiving visceral-somatic signals (Uddin et al., 2017) and the latter secreting, in the brain and in the blood circulation, neuropeptides including oxytocin associated with social bonding (Chevallier et al., 2012). In particular, the anterior part of the insula, including the right dorsal agranular region where we report positive correlation of brain activity with happiness expression by humans and negative correlation for robots’ expression (Morel et al., 2013), contains Von Economo Neurons that have a fundamental role in subjective feelings (Craig, 2009). This inversion of correlation not only signals the expected empathy for fellow humans, increase in the happiness displayed by the interlocutor triggering increase in the local response, but also the repulsion for imperfect human-like robots postulated by the Uncanny Valley hypothesis (Mori et al., 2012).

The results of the correlation analysis provide important results to understand how the regions under scrutiny are involved in social cognition. First, they support the dominance of the right hemisphere in emotional aspects of social cognition, as no significant correlation is found in the left hemisphere. In the hypothalamus, it is interesting that the brain response is similar for the two agents at the lowest levels of happiness expressed, and the difference increases with the expression of emotion, a strong argument in favour of the role of this region in the building of empathetic bonds exclusively with conspecifics, possibly through the release of oxytocin. In the insula, the results provide an interesting parcellisation with regards to social cognition. The anterior agranular regions, known to be involved in socioemotional processes, indeed provide, in the right hemisphere, the response expected for this function, as described in the previous paragraph. Dorsal regions, associated with cognitive functions, have no clear pattern in relation to social cognition. Finally, results suggest an involvement of posterior hypergranular and ventral dysgranular regions, associated with sensorimotor and chemical senses respectively (Uddin et al., 2017), in social interactions, in line with the view of embodied cognition. Differences in the patterns of these two regions are of potential interest but the absence of strong statistical significance precludes interpretations relying exclusively on social cognition explanations. For example, increased happiness by the human interlocutor could cause increased verbalization leading to a stronger reliance on sensorimotor control.

## 5 Conclusion

The analysis presented here demonstrates how robots can help to understand how the human brain processes social information during natural interactions. They demonstrate that key brain regions for emotional processing, in the framework of the somatic markers hypothesis given the choice of focusing on brain regions related to the autonomous system - insula, amygdala, and hypothalamus - respond differently to robots compared to humans’ expressions of emotions during an unconstrained conversation. Positive correlations in two key regions for social bonding, the anterior insula and hypothalamus, have their activity correlated with the level of happiness expressed by the human interlocutor. These correlations would be less interpretable if the robot control condition wasn’t available to demonstrate that these correlations are specific to human-human interactions. Our results also illustrate the grand challenge that is the development of robots capable of complex social interactions. Considering the fundamental role of emotions in human interactions, the present results directly question the acceptability of human-like robots as natural social partners. Natural interaction paradigms associated with neurophysiological recordings are particularly suited to objectively identify the strengths and weaknesses of robots in human-robot interactions. It also illustrates how crucial interdisciplinarity is for social robotics. It is with such approaches that scientists will be able to answer the crucial question: to what extent can social robots really be social?

## Data accessibility

The corpus used here is freely shared on several platforms (linguistics, behaviour, neuroscience), and details on how to access and use it will be available directly contacting the corresponding author.

## References

Adolphs, R. (2002). Neural systems for recognizing emotion. Curr Opin Neurobiol, 12(2), 169–77.

Al Moubayed, S., Beskow, J., Skantze, G., & Granström, B. (2012). Furhat: A Back-Projected Human-Like Robot Head for Multiparty Human-Machine Interaction. In A. Esposito, A. M. Esposito, A. Vinciarelli, R. Hoffmann, & V. C. Müller (Éds.), Cognitive Behavioural Systems (p. 114–130). Springer. https://doi.org/10.1007/978-3-642-34584-5_9

Arriaga, O., Valdenegro-Toro, M., & Plöger, P. G. (2019). Real-time convolutional neural networks for emotion and gender classification. ESANN 2019 - Proceedings, 27th European Symposium on Artificial Neural Networks, Computational Intelligence and Machine Learning, 221–226. https://doi.org/1710

Ashburner, J. (2007). A fast diffeomorphic image registration algorithm. Neuroimage, 38(1), 95–113. https://doi.org/10.1016/j.neuroimage.2007.07.007

Bartz, J. A., Zaki, J., Bolger, N., & Ochsner, K. N. (2011). Social effects of oxytocin in humans: Context and person matter. Trends in Cognitive Sciences, 15(7), 301–309. https://doi.org/10.1016/j.tics.2011.05.002

Belin, P. (2017). Similarities in face and voice cerebral processing. Visual Cognition, 25(4-6), 658–665. https://doi.org/10.1080/13506285.2017.1339156

Bernhardt, B. C., & Singer, T. (2012). The Neural Basis of Empathy. Annual Review of Neuroscience, 35(1), 1–23. https://doi.org/10.1146/annurev-neuro-062111-150536

Blouin, A. M., Fried, I., Wilson, C. L., Staba, R. J., Behnke, E. J., Lam, H. A., Maidment, N. T., Karlsson, K. Æ., Lapierre, J. L., & Siegel, J. M. (2013). Human hypocretin and melanin-concentrating hormone levels are linked to emotion and social interaction. Nature Communications, 4(1), 1547. https://doi.org/10.1038/ncomms2461

Brett, M., Anton, J. L., Valabregue, R., & Poline, J. B. (2002). Region of interest analysis using an SPM toolbox. 8th international conference on functional mapping of the human brain, 16, 497. http://dx.doi.org/10.1016/S1053-8119(02)90010-8

Chaminade, T. (2017). An experimental approach to study the physiology of natural social interactions. Interaction Studies, 18(2), 254–275.

Chaminade, T., Da Fonseca, D., Rosset, D., Cheng, G., & Deruelle, C. (2015). Atypical modulation of hypothalamic activity by social context in ASD. Research in Autism Spectrum Disorders, 10, 41–50.

Chaminade, T., Zecca, M., Blakemore, S.-J., Takanishi, A., Frith, C. D., Micera, S., Dario, P., Rizzolatti, G., Gallese, V., & Umiltà, M. A. (2010). Brain response to a humanoid robot in areas implicated in the perception of human emotional gestures. PLoS one, 5(7), e11577.

Chevallier, C., Kohls, G., Troiani, V., Brodkin, E. S., & Schultz, R. T. (2012). The social motivation theory of autism. Trends in Cognitive Sciences, 16(4), 231–239. https://doi.org/10.1016/j.tics.2012.02.007

Craig, A. D. (2009). How do you feel—Now ? The anterior insula and human awareness. Nature Reviews Neuroscience, 10(1), 59–70. https://doi.org/10.1038/nrn2555

Crozier, W. R. (2001). Blushing and the exposed self: Darwin revisited. J Theroy soc Behav, 31(1), 61–72.

Damasio, A. R., Everitt, B. J., & Bishop, D. (1996). The Somatic Marker Hypothesis and the Possible Functions of the Prefrontal Cortex [and Discussion]. Philosophical Transactions: Biological Sciences, 351(1346,), 1413–1420.

Damasio, A. R., Grabowski, T. J., Bechara, A., Damasio, H., Ponto, L. L., Parvizi, J., & Hichwa, R. D. (2000). Subcortical and cortical brain activity during the feeling of self-generated emotions. Nat Neurosci, 3(10), 1049–56.

Darwin, C. (1872). The Expression of the Emotions in Man and Animals by Charles Darwin. Murray.

Ekman, P. (1974). Darwin and Facial Expression: A Century of Research in Review. Academic Press Inc.

Fan, L., Li, H., Zhuo, J., Zhang, Y., Wang, J., Chen, L., Yang, Z., Chu, C., Xie, S., Laird, A. R., Fox, P. T., Eickhoff, S. B., Yu, C., & Jiang, T. (2016). The Human Brainnetome Atlas: A New Brain Atlas Based on Connectional Architecture. Cereb Cortex, 26(8), 3508–3526. https://doi.org/10.1093/cercor/bhw157

Friston, K. J., Ashburner, J. T., Kiebel, S., Nichols, T. E., & Penny, W. D. (2007). Statistical Parametric Mapping: The Analysis of Functional Brain Images. Elsevier.

Harrison, N. A., Gray, M. A., & Critchley, H. D. (2009). Dynamic pupillary exchange engages brain regions encoding social salience. Social Neuroscience, 4(3), 233–243. https://doi.org/10.1080/17470910802553508

Hortensius, R., Hekele, F., & Cross, E. S. (2018). The Perception of Emotion in Artificial Agents. IEEE Transactions on Cognitive and Developmental Systems, 10(4), 852–864. https://doi.org/10.1109/TCDS.2018.2826921

Kasper, L., Bollmann, S., Diaconescu, A. O., Hutton, C., Heinzle, J., Iglesias, S., Hauser, T. U., Sebold, M., Manjaly, Z.-M., Pruessmann, K. P., & Stephan, K. E. (2017). The PhysIO Toolbox for Modeling Physiological Noise in fMRI Data. Journal of Neuroscience Methods, 276, 56–72. https://doi.org/10.1016/j.jneumeth.2016.10.019

LeDoux, J. (2007). The amygdala. Current Biology: CB, 17(20), R868–874. https://doi.org/10.1016/j.cub.2007.08.005

Morel, A., Gallay, M. N., Baechler, A., Wyss, M., & Gallay, D. S. (2013). The human insula: Architectonic organization and postmortem MRI registration. Neuroscience, 236, 117–135. https://doi.org/10.1016/j.neuroscience.2012.12.076

Mori, M., MacDorman, K. F., & Kageki, N. (2012). The Uncanny Valley [From the Field]. IEEE Robotics Automation Magazine, 19(2), 98–100. https://doi.org/10.1109/MRA.2012.2192811

Pessoa, L. (2017). A Network Model of the Emotional Brain. Trends in Cognitive Sciences, 21(5), 357–371. https://doi.org/10.1016/j.tics.2017.03.002

Poppa, T., & Bechara, A. (2018). The somatic marker hypothesis: Revisiting the role of the ‘body-loop’ in decision-making. Current Opinion in Behavioral Sciences, 19, 61–66. https://doi.org/10.1016/j.cobeha.2017.10.007

Rauchbauer, B., Hmamouche, Y., Bigi, B., Prevot, L., Ochs, M., & Thierry, C. (2020). Multimodal corpus of bidirectional conversation of human-human and human-robot interaction during fMRI scanning. Proceedings of The 12th Language Resources and Evaluation Conference, 661–668.

Rauchbauer, B., Nazarian, B., Bourhis, M., Ochs, M., Prévot, L., & Chaminade, T. (2019). Brain activity during reciprocal social interaction investigated using conversational robots as control condition. Philosophical Transactions of the Royal Society B, 374(1771), 20180033.

Redcay, E., & Schilbach, L. (2019). Using second-person neuroscience to elucidate the mechanisms of social interaction. Nature Reviews Neuroscience, 20(8), 495–505. https://doi.org/10.1038/s41583-019-0179-4

Saygin, Z. M., Osher, D. E., Augustinack, J., Fischl, B., & Gabrieli, J. D. E. (2011). Connectivity-based segmentation of human amygdala nuclei using probabilistic tractography. NeuroImage, 56(3), 1353–1361. https://doi.org/10.1016/j.neuroimage.2011.03.006

Uddin, L. Q., Nomi, J. S., Hébert-Seropian, B., Ghaziri, J., & Boucher, O. (2017). Structure and Function of the Human Insula. Journal of Clinical Neurophysiology: Official Publication of the American Electroencephalographic Society, 34(4), 300–306. https://doi.org/10.1097/WNP.0000000000000377

Vuilleumier, P., & Pourtois, G. (2007). Distributed and interactive brain mechanisms during emotion face perception: Evidence from functional neuroimaging. Neuropsychologia, 45(1), 174–194. https://doi.org/10.1016/j.neuropsychologia.2006.06.003

Wolfe, F. H., Auzias, G., Deruelle, C., & Chaminade, T. (2015). Focal atrophy of the hypothalamus associated with third ventricle enlargement in autism spectrum disorder. Neuroreport, 26(17), 1017–1022.

